# The role of carbonic anhydrase in rock weathering and pH regulation by the soil bacterium *Burkholderia thailandensis* E264

**DOI:** 10.1101/2024.11.21.624755

**Authors:** Derek S. Bell, Jonathan R. Leake, David J. Beerling, Jurriaan Ton, Dimitar Z. Epihov

## Abstract

Enhanced rock weathering (ERW) is increasingly recognized as a way to sequester atmospheric carbon dioxide (CO^2^) to slow global warming, but its effectiveness needs to be optimized. Carbonic anhydrase (CA), an enzyme capable of accelerating rock weathering both *in vitro* and in soil, offers a valuable target due to its ability to convert CO^2^ into carbonic acid. This conversion promotes rock dissolution, enabling immediate CO^2^ absorption through cation release and charge-balance mechanisms. Studies have shown that bacteria grown in axenic, rock-amended media increase CA gene expression, but the influence of bacterial CA on rock dissolution rates remains unclear. To investigate this, we used a reverse genetics approach with the phosphate-solubilizing bacterium *Burkholderia thailandensis* E264. We examined three CA-inactivated mutants alongside the wildtype, growing them in minimal media with basalt rock dust (0-10% w/v) at an initial pH of 6.0. After 7 days, we measured weathering potential through elemental concentrations, pH, and dissolved inorganic carbon. In 1% basalt medium, inactivation of the CA1 gene (BTH_I1052) significantly reduced base cation weathering by 41% compared to the wildtype, whereas inactivation of CA2 (BTH_I0345) and CA3 (BTH_I1199) had no significant effect. In the highly buffered, 10% basalt medium, CA1 had a minor role in weathering, and both CA2 and CA3 had no effect. These findings suggest that CA genes in *B. thailandensis* operate differently and that CA1’s effect is pH-dependent. Surprisingly, CA1 was localized intracellularly, raising questions about how intracellular CAs might influence mineral dissolution, potentially through acidity export or abiontic enzyme activity after cell lysis.

**Importance:** While purified carbonic anhydrase (CA) protein has been shown to increase mineral dissolution rates in mineral-amended media *in vitro*, it remains unclear if the bacterial CA gene directly drives this process. This study used CA-inactivated mutants of the soil bacterium *Burkholderia thailandensis* in basalt-amended liquid media and found that only one of the three CA genes influenced mineral dissolution rates. This finding supports prior evidence that bacterial CAs may contribute to mineral dissolution in soils. Importantly, it also showed that not all CA genes in a bacterium may activate under the same conditions, which could impact how soil bacterial CAs are leveraged to enhance weathering. Furthermore, cellular localisation predictions indicated that all three CA genes in *B. thailandensis* are cytosolic, challenging the common focus on extracellular CAs and suggesting that CA proteins may influence the external environment without needing to be actively exported from the cell.

## 1. Introduction

Rock weathering is a geochemical process by which rocks physically and chemically break down due to the action of the atmosphere, water, and/or organisms, driving long-term carbon sequestration within the inorganic carbon cycle, that regulates Earths atmospheric CO_2_ concentration and climate over geological timescales (Berner, 2004; Taylor et al., 2009). When carbonic acid reacts with calcium or magnesium silicates it results in carbon dioxide (CO_2_) sequestration into bicarbonate or minerals such as calcium carbonate. This naturally occurring process is currently too slow to balance additional anthropogenic CO_2_ emissions (Hartmann et al., 2009). Carbonic acid formation, a product of CO_2_ dissolution in water, can be catalysed by carbonic anhydrase (CA) (Figure 1). This weak acid can then react with calcium or magnesium silicates to draw down 2 moles of CO_2_ in the form of bicarbonate.

**Figure 1.** Wollastonite weathering reactions, with CA activity highlighted.

Enhanced rock weathering (ERW) is a land-based CO_2_ removal (CDR) strategy, involving the spreading of crushed calcium-bearing silicate rock onto agricultural soils to artificially increase rates of calcium and magnesium silicate weathering (Schuiling and Krijgsman, 2006; Moosdorf et al., 2011; Hartmann et al., 2013). Global-scale modelling suggests that if ERW is rolled out over ∼50% of croplands for major nations of the world it could result in a net sequestration of 0.5 – 2.0 Gt CO_2_ yr^-1^ by 2050 (Beerling et al., 2020). In addition to the benefits of CDR, numerous short-term field trials reviewed by Abdalqadir et al. (2024), along with the first long-term ERW field trial conducted by Beerling et al. (2024), have shown that basaltic rock amendments to soils significantly increased soil pH, inorganic nutrient concentrations, and crop yields.

Finding methods to accelerate rates of ERW within agricultural soils is important to optimize CDR rates (Epihov et al., 2024) and poses unique challenges due to the complexity of the soil-mineral-enriched environment. However, Ribeiro et al. (2020) reviewed numerous bacterially driven biological weathering mechanisms which could potentially be artificially enhanced. The catalysis of the formation and dissolution of carbonic acid in water by CA (Figure 1) provides a potentially important target for such enhancement. Soil respiration, deriving from microbiota, plant roots, and fauna leads to high concentrations of CO_2_ within the soil solution, which naturally drives the formation of carbonic acid with CA estimated to increase the rate of carbonic acid formation in the soil by 10 – 300 times (Wingate et al., 2009). Consequently, there is considerable interest in understanding its potential role in accelerating ERW with bacteria (Li et al., 2007, 2009; Dhami et al., 2014; Xiao et al., 2015; Shen et al., 2017; Wang et al., 2018; Jaya et al., 2019; Nathan and Ammini, 2019) and fungi (Liu and Dreybrodt, 1997; Han et al., 2010; Xiao et al., 2016; Sun and Lian, 2019).

Current research into CA generally focuses on the human α-CA isoforms or pathogenic organisms (Nocentini and Supuran, 2019). This research provides useful information regarding the catalytic efficiencies of different CAs, with β-CA’s varying up to 10-fold between different organisms (Table 1). Similar variations are also noted between the 16 different α-CA isoforms present in mammals (Pinard et al., 2015). These variations in catalytic efficiencies are likely due to structural differences between members within both the α and the β-CA classes. This indicates that there is unlikely to be a single CA activity profile in different soils, but instead large variations depending on which microorganisms are prevalent in the soil.

**Table 1.** β-CA catalytic potentials in bacteria 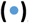, fungi 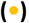, archaea 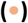, eukaryotes 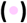, and plants 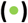. N/A not available.

| Organism | Enzyme | Kcat ($s^{-1}$) | Kcat per Km ($M^{-1}s^{-1}$) | pH | Source |
| --- | --- | --- | --- | --- | --- |
| *Pseudomonas aeruginosa* | psCA1 | $1.8 \times 10^5$ | $7.5 \times 10^7$ | 8.3 | (Lotlikar et al., 2019) |
| *Helicobacter pylori* | hpβCA | $7.1 \times 10^5$ | $4.8 \times 10^7$ | 8.3 | (Nishimori et al., 2007) |
| *Escherichia coli* | CynT2 | $5.3 \times 10^5$ | $4.1 \times 10^7$ | 8.3 | (Del Prete et al., 2020) |
| *Vibrio cholerae* | VchCAβ | $3.3 \times 10^5$ | $4.1 \times 10^7$ | 8.3 | (Ferraroni et al., 2015) |
| *Mycobacterium tuberculosis* | mtCA 1 | $3.9 \times 10^5$ | $3.7 \times 10^7$ | 8.3 | (Minakuchi et al., 2009) |
| *Burkholderia pseudomallei* | BpsβCA | $1.6 \times 10^5$ | $3.4 \times 10^7$ | 8.3 | (Angeli et al., 2020) |
| *Saccharomyces cerevisiae* | scCA | $9.4 \times 10^5$ | $9.8 \times 10^7$ | 8.3 | (Isik et al., 2008) |
| *Cryptococcus neoformans* | Can2 | $3.9 \times 10^5$ | $4.3 \times 10^7$ | 8.3 | (Innocenti et al., 2008) |
| *Candida albicans* | Nce103 | $8.0 \times 10^5$ | $9.7 \times 10^7$ | 8.3 | |
| *Methanobacterium thermoautotrophicum* | Cab | $1.7 \times 10^4$ | $5.9 \times 10^6$ | 8.5 | (Smith and Ferry, 1999) |
| *Leishmania donovani chagasi* | LdcCA | $9.4 \times 10^5$ | $5.9 \times 10^7$ | 8.4 | (Syrjä et al., 2013) |
| *Pisum sativum* | N/A | $1 \times 10^6$ | $1.8 \times 10^9$ | N/A | (Kimber, 2000) |
| *Flaveria bidentis* | FbiCA 1 | $1.2 \times 10^5$ | $7.5 \times 10^6$ | 8.4 | (Monti et al., 2013) |

The first evidence showing CA’s potential role in rock weathering was presented by Liu and Dreybrodt (1997), in which they noted that the addition of bovine α-CA increased the rate of limestone dissolution in an *in vitro* system. Subsequently, it was shown that bacteria can induce weathering within an axenic soil medium containing limestone, as their addition increased the concentration of calcium in the soil leachate (Li et al., 2005). When a known CA inhibitor, acetazolamide, was added to a bacterial or crude enzyme mixture there was less calcium released from the limestone over the course of 9 hours (Li et al., 2007). More recent research has examined the genetic regulation of CA and attempts have been made to purify the enzyme (Xiao et al., 2015; Supuran, 2013). Xiao et al. (2015) presented promising results showing that all 5 CA encoding genes within *Bacillus mucilaginosus* were significantly upregulated following six days of growth in a wollastonite amended medium. They also showed that a purified bacterial CA protein (PCA4) when incubated at 0.039% CO_2_ increased the rate of wollastonite dissolution by 4.78 × 10^−4^ mg g^-1^ min^-1^. This evidence strongly suggests that CA plays a role in the weathering of rocks, but another *in vitro* study suggests a more complex dynamic as CA inhibited rock weathering (Di Lorenzo et al., 2018). This contradiction might be due to differences in experimental conditions since Di Lorenzo et al. (2018) ran their experiment with a pH fixed to 4; whereas Xiao et al. (2015) began the reaction at a pH of approximately 8 and allowed it to change. Through measuring ^18^O enriched CO_2_ flux of soils ranging from pH 4.5 – 8.5 with bovine α-CA added to the soils, Sauze et al. (2018) demonstrated that soil pH is the primary determinant of soils CO_2_ – H_2_O isotopic exchange rate. Their results indicate that at a low soil pH, even with addition of CA, rates of CO_2_ hydration did not change compared to the native control; whereas, at a higher soil pH CA addition increased the rate of CO_2_ hydration by 20 reactions sec^-1^ (Sauze et al., 2018). Since calcium and magnesium-rich rock dusts tend to increase soil alkalinity, this suggests that CA may be an important facilitator of ERW. Research into fungal CA in *Aspergillus nidulans* has further indicated its active role in mineral weathering; when grown in a wollastonite amended minimal medium with the CA gene *can*A inactivated, there was a loss of soluble calcium in the medium, whereas overexpression of *can*A resulted in an increase in soluble calcium (Sun and Lian, 2019). All the aforementioned literature has generated commercial interest, leading to a partnership between the two companies, Veolia and FabricNano, which has initiated field trials by adding a CA inoculum to soils, both freely and immobilized on rock surfaces (Peplow, 2024).

The cellular localization of the CA may also play a role in its effectiveness as a rock weathering enzyme; Figure 2 summarises multiple ways CA could affect its extracellular environment. Experiments with bovine α-CA added have assumed that the enzyme is most active extracellularly. In addition, four sequences obtained for the CAs examined by Xiao et al., (2015) were also estimated as having extracellular localisations through testing with PSORTb v. 3.0.3 (Yu et al., 2010).

**Figure 2.**
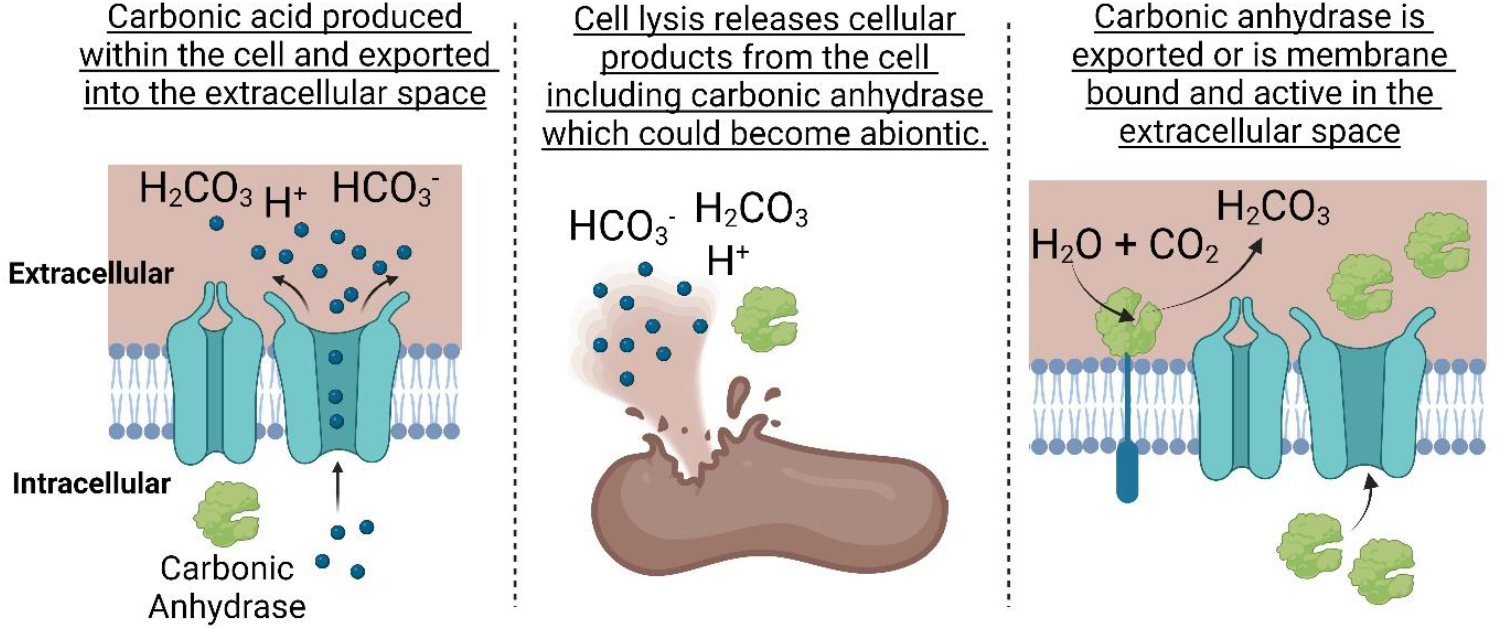
Potential CA methods of extracellular chemical change. Created in BioRender.com

More recent *in vitro* ERW research has used mutants from the transposon-mutant library of the soil β-proteobacterium *Burkholderia thailandensis* E264, and has seen that CA was significantly correlated with respiration genes in basalt amended tropical soils (Epihov, 2018). *B. thailandensis*, a gram negative, aerobic, motile, and non-fermenting bacterium, is a widespread soil saprophyte in Australasia (Zakharova et al., 2022). It is also commonly used as a model organism to study the highly pathogenic *Burkholderia pseudomallei*, which has resulted in it having a well characterised genome along with the aforementioned mutant collection (Gallagher et al., 2013). *B. thailandensis* is also considered to be a plant growth promoting bacterium as when it was applied to rice growing in acid sulfate soils, along with a basalt and magnesium limestone, it reduced the rhizotoxicity of aluminium and increased solubilised phosphate and yields (Panhwar et al., 2015). Experiments have confirmed its plant growth promoting potential through identifying not only phosphate solubilization but also nitrogen fixation genes (Sivaji et al., 2016). It can also inhibit the growth of the commercial crop pathogen *Rhizoctonia solani* (Sivaji et al., 2016), and can degrade the insecticide Fipronil (Cappelini et al., 2018). Its potential importance as a plant growth promoting bacterium in cropping systems and its well-characterised genome makes it a suitable candidate for the present study. Our study aims to use a reverse genetics approach through growing the *B. thailandensis* wildtype strain and CA inactivated mutants in media amended with different concentrations of basalt rock to assess the role of bacterial CA in rock dissolution and pH regulation.

## 2. Materials & Methods

### 2.1 Mineral

Basalt, sourced from The Hillhouse Quarry Group in Troon, Ayrshire, Scotland, was composed of 34.6% Plagioclase, 32.7% augite, 14.7% forsterite, 5.2% chlorite-smectite, 4.5% spinel, 7.0% analcime, and 0.5% Ilmenite (Lewis et al., 2021). The rock was first dry sieved to sizes of 75 - 90 µm and < 53 µm, the 75 - 90 µm fraction underwent additional washing with 0.1 M HCl to remove exchangeable bases (Yu et al., 2016) followed by rinsing with distilled water until all visible dust particles were removed (Epihov, 2018). Both fractions were dried at 100 °C overnight, with 0.35 g of the 75 - 90 µm (1% w/v basalt medium) or 3.5 g of the < 53 µm size fraction (10% w/v basalt medium) placed in autoclavable 50 mL falcon tubes. These tubes were loosely sealed, placed in an autoclavable bag, and wet autoclaved at 121 °C for 45 min twice, with at least 24 hours at room temperature between autoclave cycles.

### 2.2 Bacteria preparation

The wildtype (WT) strain E264 and a knockout (KO) library of the soil bacterium *Burkholderia thailandensis* were acquired from the Manoil lab at the University of Washington, U.S (Gallagher et al., 2013). The KO library included three mutants that had CA genes inactivated, the loci included: BTH_I1052, BTH_I0345, and BTH_I1199, referred to in the following text as CA1, CA2, and CA3, respectively. Colonies of both WT and KO organisms were grown from - 80 °C stocks on streak plates incubated overnight at 37 °C on Luria Bertani (LB) agar (Bertani, 1951). Single colonies from each respective organism were used to inoculate 30 mL LB broths, which were grown aerobically with orbital shaking at 250 rpm (New Brunswick Innova® 44/44R) overnight at 37 °C along with a non-inoculated control. Following growth, optical density at 600 nm (Geneva Bio, Jenway) were taken and broths with higher values were diluted using sterile LB broth to match the lowest absorbance. All tubes were then centrifuged at 3234 x *g* for 10 minutes and supernatants were aseptically removed prior to the pellets being resuspended in 10 mL of 0.85% sterile isotonic saline. This washing step was repeated two more times, with the final pellet being resuspended in 12 mL of 0.85% sterile saline before inoculation into the media, described in section 2.3.

### 2.3 Media preparation, inoculation, and growth

The minimal media, composed of 10 mM D-glucose, 1 mM sodium chloride, 0.020 mM magnesium sulfate heptahydrate, 0.050 mM dipotassium phosphate, and 1 mM ammonium nitrate, was prepared in 1 L of distilled ultra-pure water. Using the previously weighed and autoclaved rock described in section 2.1, three replicates with concentrations of 0, 1, and 10% w/v basalt were created for all four organisms described in section 2.2, alongside with six non-inoculated controls. This process involved adding 34.9, 34.775, and 33.65 mL of sterilized minimal media to tubes containing 0, 0.35, and 3.5 g of basalt rock, respectively.

The bacterial suspensions were thoroughly vortexed before adding 100 µL of each relevant organism or non-inoculated control to their respective 50 mL Falcon tubes, containing either 0, 1 or 10% w/v basalt. Following inoculation, a colony forming units (CFU) dilution series was performed on LB agar and counted after overnight incubation aerobically at 37 °C for each organism and the control to determine initial numbers of bacteria inoculated.

Three of the non-inoculated controls of each basalt concentration were sampled to determine initial pH of the media. The remaining samples were incubated aerobically with 120 rpm shaking at 37 °C for 170 hours. CFU dilution series and counts were performed post-inoculation at time points: 3, 6, 9, 21, 24, 27, 48, and 170 hours. Total growth was determined using CFU counts because optical density has been found to be an unreliable method of growth with *B. thailandensis* due to its secreted metabolites (Martinez and Déziel, 2020).

### 2.4 Sample analysis and CO_2_ absorption estimate

At 170 hours, all tubes were centrifuged at 3234 x *g* for 20 minutes at 4 °C. The resulting supernatants were filter sterilised to 0.2 µm and transferred into 50 mL Falcon tubes. These samples were kept on ice, with aliquots removed for further analyses. For ICP-MS analysis, 1.2 mL of each sample was diluted with a mixture of 10.452 mL ultra-pure water and 0.348 mL of 69% nitric acid, resulting in a 1/10 diluted sample with 2% nitric acid. Subsequently, all tubes were sent for ICP-MS analysis with the Agricultural and Environmental Sciences Department at the University of Nottingham. Total inorganic carbon analysis was conducted by removing 9 mL of the filtered sample and placing in a sample test tube tightly sealed with polyethylene cling film to prevent CO_2_ degassing (Sun and Li, 2017) until all samples could be run on the TOC-L (Shimadzu). The final pH of all experimental and control tubes was measured using the Bante930 Benchtop pH/Ion Meter equipped with an P11 glass electrode calibrated to pH’s 4.00, 7.00, and 10.00.

Potential CDR was estimated using a modified technique of charge balancing base cations calcium and magnesium with bicarbonate ions, as described previously in soils (Hartmann et al., 2013; Kanzaki et al., 2023; Beerling et al., 2024). Briefly, the net molar change of cations calcium and magnesium in the media due to basalt amendment was totalled and using a 2:1 molar ratio of HCO_3_^-1^ to cations, the CO^2^ sequestration in g L^-1^ was estimated.

### 2.5 Statistical analysis

GraphPad Prism version 10.0.0 for Windows (GraphPad Software, Boston, Massachusetts USA, www.graphpad.com) was used for plotting all figures. Normality and homoscedasticity were assessed visually using residual and Q-Q plots available within R studio (R Core Team, 2024). Using the Mass package in R (Venables and Ripley, 2002) data with non-normal distribution of residuals underwent boxcox analysis to identify the optimal lambda transformation to normalise the residuals (Box and Cox, 1964). Using Core R packages both two-way and three-way ANOVAs and Tukey multiple comparisons statistical tests were also performed in R. A result was considered significant if the *p* value was < 0.05.

## 3. Results

### 3.1 Total growth, final pH, and dissolved inorganic carbon

Prior to inoculation, the average initial pH values of the 0, 1 and 10% w/v basalt media were 6.55, 6.51, and 8.71, respectively. The higher initial pH of the 10% w/v media was expected due to the presence of < 53 µm rock grains, which likely increased the concentration of dissolved base cations in the solution. However, after 170 hours of incubation at 37 °C, the average pH of the uninoculated control samples containing 10% w/v basalt fell to 7.52. Bacterial growth in the basalt-free medium caused consistent acidification, whereas in the 1% w/v basalt media, acidification was less pronounced and showed more variability between replicates and organisms. For example, the KO CA1 appeared distinctive as it maintained a higher average pH of 7.1 compared to the wildtype which had a pH of 5.1. Post-hoc Tukey’s tests confirmed a significantly higher pH for this mutant compared to the other organisms (Figure 3a).

**Figure 3.**
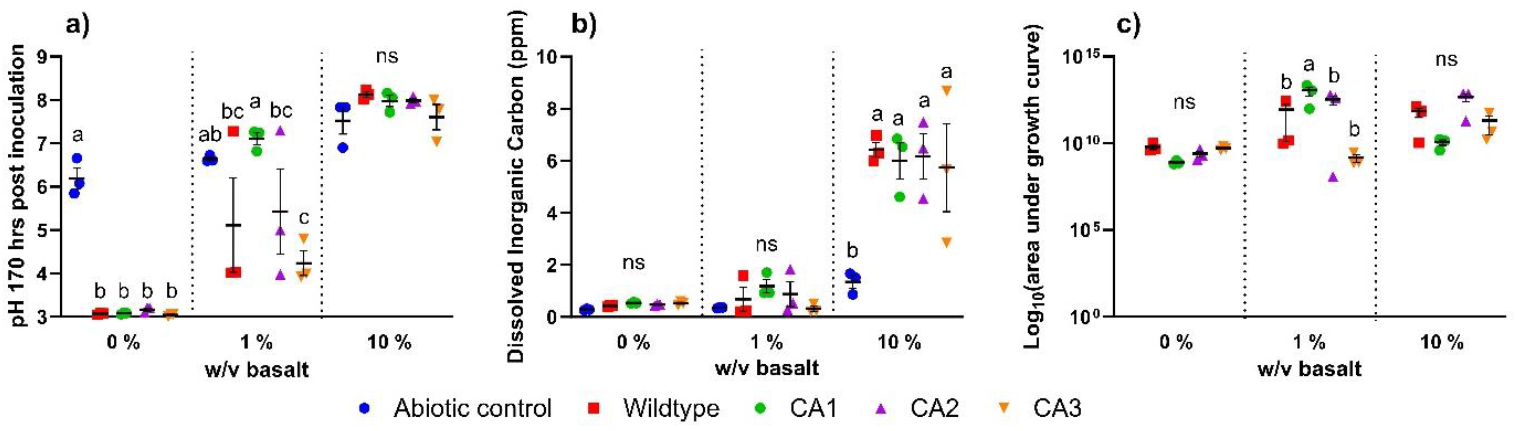
Effects of basalt treatments and *B. thailandensis* growth 170 hours post inoculation compared to abiotic controls on mean (± 1 standard error of mean, SEM) (a) pH, (b) dissolved inorganic carbon, and (c) total area under growth curve. ANOVA post hoc Tukey’s test significance differences are denoted by letter codes, means sharing the same letter are not significantly different (*p* > 0.05) and “ns” denotes no significant pairwise differences.

The dissolved inorganic carbon found in the media post incubation (Figure 3b), showed no significant pairwise differences between organisms at any basalt concentration. The total growth over the course of the experiment, (Figure 3c) expressed as log_10_(area under the growth curve), showed a significant increase in the growth of the CA1 KO when grown with 1% w/v basalt relative to the other organisms, but there were no differences between organisms in the 0% and 10% basalt treatments. Importantly, no bacterial growth was seen in any of the abiotic controls.

### 3.2 Rock dissolution and bacterial enhanced element release into solution

When grown without basalt, elemental concentrations relative to the control (Figure 4) remained consistently low for all organisms, except for the CA2 mutant, which showed significant increases in potassium, selenium, rubidium, silver, and caesium. As seen in Figure 4, inoculation of *B. thailandensis* into basalt amended medias in most cases increased the total dissolved elemental concentrations (DEC), indicating that bacteria increased weathering rates in the media. Interestingly, in the 1% w/v basalt media (Figure 4b) the KO CA1 behaved differently to the other organisms, with no elements significantly increasing relative to the control. However, in both the 1% and 10% w/v media the CA1 mutant had significantly reduced concentrations of phosphorus and silver. Along with those reductions in the 10% media the CA1 mutant also had significant increases in the concentrations of calcium, magnesium, strontium, rubidium, and uranium but these significant increases were also observed with the other KOs and the wildtype.

**Figure 4.**
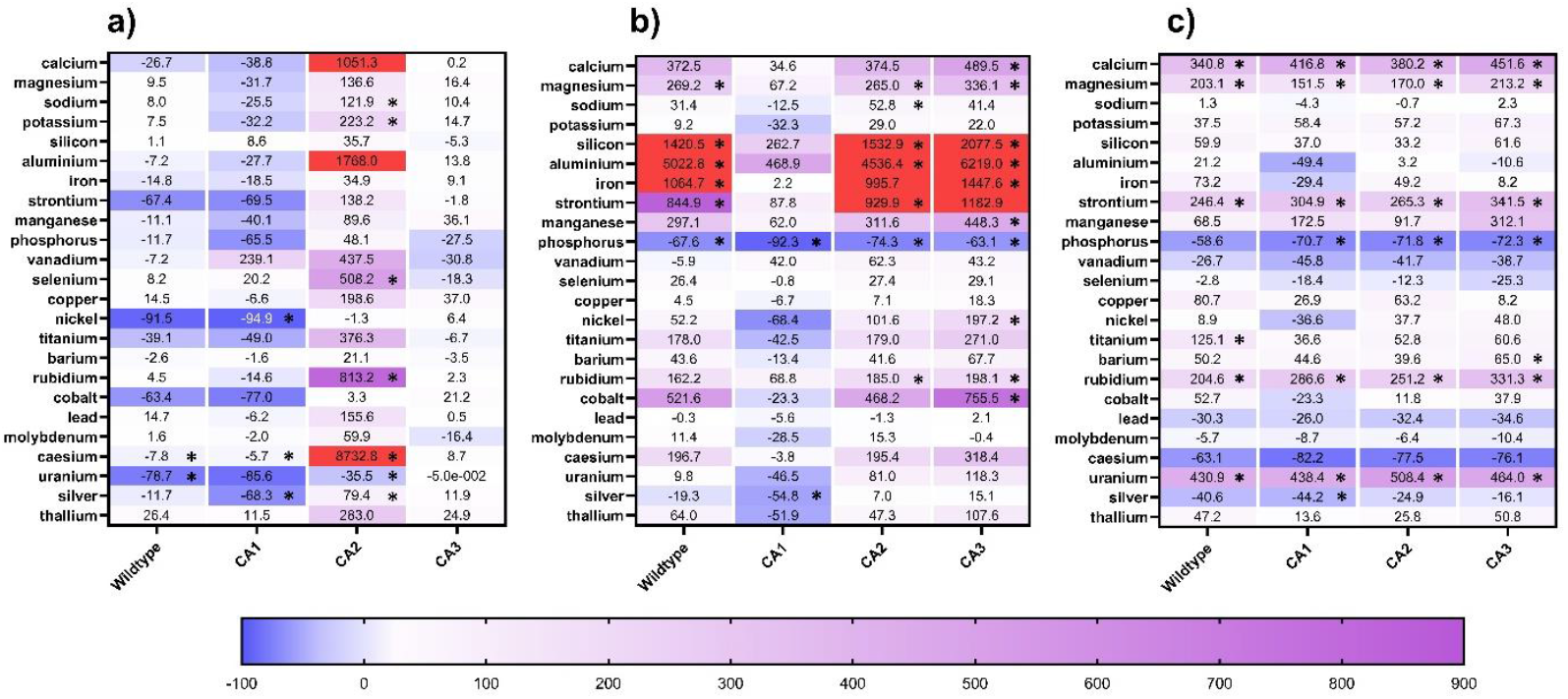
Heat map showing average dissolved element percent change relative to the abiotic control 170 hours post inoculation of *B. thailandensis* organisms wildtype, CA1, CA2, and CA3 in media with (a) 0, (b) 1%, and (c) 10% w/v basalt. Red cells indicate values exceeding the heat map range. Cells that are significantly different (*p* values < 0.05) to the control are denoted by an asterisk (*).

Supplemental Table 1 shows all measured elements that could be statistically evaluated using a two-way ANOVA and post-hoc Tukey’s test. As expected in the 0% w/v media, DECs tended to be lower than the basalt amended medias. Furthermore, when the KO CA2 was grown in the 0% w/v media it resulted in significantly greater DECs of sodium, selenium, copper, and rubidium relative to the other organisms. This trend was observed for other elements, however it was not significant.

When grown in 1% w/v media, a pattern was seen across almost all the elements, excluding vanadium. Specifically, CA1 consistently had a lower mean DEC relative to the other KOs and the wildtype. This was significantly different from all other organisms for strontium (80.12% less, *p* < 0.05) and silver (99.64% less, *p* < 0.05) relative to the wildtype. For the rest of the elements measured, this trend of CA1 giving lower concentrations showed varying significant differences compared to the other organisms. The only element that contradicted this trend was vanadium, where CA1’s mean concentration was 0.19 µg L^-1^ greater than the wildtype, though this increase was not significant (*p* > 0.05).

In the 10% basalt media *B. thailandensis* colonisation significantly increased cation concentrations of calcium, magnesium, and potassium (Supplemental Figure 1 a, b, d), with no significant differences noted between organisms. Excluding caesium, all other elements showed no significant differences between organisms; however, a pattern was seen in DEC for the elements titanium, vanadium, aluminium, cobalt, nickel, silver, caesium, iron, and barium suggesting that these dissolved elements may be released at lower rates due to CA inactivation. When titanium, vanadium, cobalt, nickel, silver, and caesium are averaged there is a significant three-way ANOVA difference (*p* < 0.001) between the wildtype and the CA inactivated mutants. However, only the wildtype and CA3 were significantly greater (*p* < 0.001) than the CA1 KO by 46 and 48 %, respectively.

Interestingly, some elements, such as phosphorus, vanadium, and selenium, appeared to decrease because of *B. thailandensis* growth. Of those elements, phosphorus was the only one to see significant reductions relative to the control of 0.127 (*p* < 0.05), 0.124 (*p* < 0.05), and 0.126 (*p* < 0.05) mg L^-1^ when the *B. thailandensis* KO organisms CA1, CA2, and CA3 were grown in the 10% w/v media, respectively.

When all measured elemental concentrations presented in Supplemental Figure 1 were summed, averaged across replicates, and analysed using a two-way ANOVA with post-hoc Tukey’s test (Figure 5a), similar patterns were observed to those in Figure 4 and Supplemental Figure 1. Specifically, the total averaged DEC in the 0% w/v media showed a greater mean DEC when the CA2 KO was grown, but when elements were totalled this became non-significant. In the 1% w/v media, the CA1 KO organism had a significant reduction in DECs relative to the other KOs, but not to the control or the wildtype. No significant pairwise differences were observed in total DEC’s between organisms grown in the 10% w/v media, but all organisms differed significantly from the control, again highlighting the importance of bacteria in rock dissolution. When the cations calcium, magnesium, sodium, and potassium were totalled (Figure 5b, Supplemental Figure 1), the distribution of means was similar to that in Figure 5a at all basalt concentrations. However, in this case, the CA2 KO had significantly higher concentrations than both the control and the other organisms for these elements when grown in 0% w/v basalt, while the CA1 KO, when grown in 1% w/v basalt, had significantly reduced dissolved cation concentrations relative to all other organisms, including the wildtype. In the 10% w/v basalt media, total dissolved cations appeared to increase relative to the control, but only the CA3 KO was significantly different. Estimated potential CO_2_ absorption (Figure 5c), calculated as described in Section 2.4, revealed a distribution of means similar to those in Figures 5a and 5b. However, as in Figure 5b, the CA2 KO, when grown in 0% w/v media, was significantly different from the control. Additionally, when grown in 1% w/v media, the CA1 KO was significantly reduced relative to the other KOs but not different from the wildtype; in this case, though, the wildtype was different from the control.

**Figure 5.**
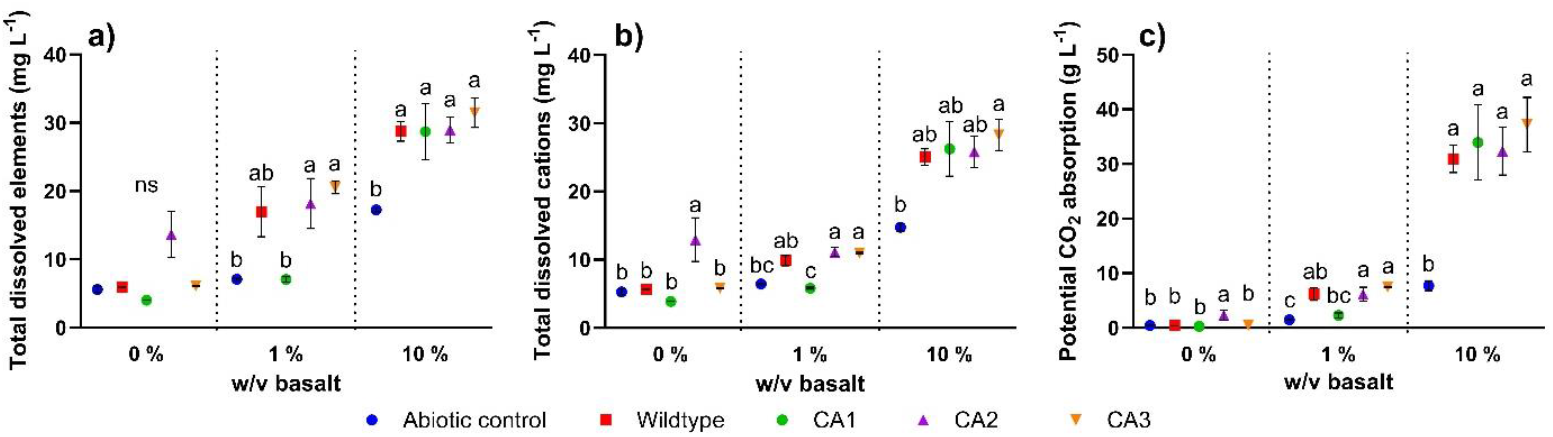
Effects of basalt amendment and *B. thailandensis* wildtype and mutant growth, along with abiotic controls 170 hours post-inoculation, on mean concentrations (± 1 standard error) of (a) total dissolved elements (b) total dissolved cations (calcium, magnesium, sodium, and potassium), and (c) potential CO_2_ absorption. Two-way ANOVA post hoc Tukey’s test significance differences denoted by letters codes within basalt treatments, where means sharing the same letter are not significantly different (*p* > 0.05) and “ns” denotes no significant pairwise differences.

## 4. Discussion

Previous research has suggested that *Bacillus subtilis* inoculation *in vitro* and *in vivo* could increase calcium liberation from silicate rock to facilitate ERW (Timmermann-Aranis et al., 2024). However, this may not necessarily increase carbonic acid derived weathering. CA, however, is the only proposed mechanism which should theoretically directly increase CDR rates in ERW, as it catalyses the formation of carbonic acid. Currently, two commercial companies (FabricNano and Veolia) are trialling a mineral immobilised CA enzyme (Peplow, 2024) for increasing weathering rates, with the idea that the enzyme is stabilised in the alkaline environment of the “mineralosphere” (Uroz et al., 2015) optimal for CA activity. Another potential method of increasing CDR rates with CA is to leverage the CA’s within the soil community, but more research is needed to characterise the role of CA within soil bacteria, and under what circumstances the enzyme could increase weathering rates. We addressed this issue through using a reverse genetics approach to characterise CA’s role in *in vitro* rock weathering under a weakly buffered and highly buffered rock amended media.

We show that in the weakly buffered 1% w/v basalt media, the inactivation of the CA1 gene appeared to play more of a role in rock weathering; as it released significantly fewer total cations into the solution, despite achieving more cell growth than the other organisms. This trend of reduced DECs with the KO CA1 was noted for all measured elements. When totalled across elements and averaged, the inactivation of the CA1 gene resulted in significantly lower element concentrations in solution relative to the CA2 (61.2% less, *p* < 0.05) and CA3 (65.6% less, *p* < 0.05) mutants and a lower, albeit non-significant, reduction relative to the wildtype (58.4% less, *p* > 0.05). This diminished weathering potential occurred along with it carrying a final pH average of 7.11, which was significantly higher than the other organisms. The elevated pH seen with the CA1 mutant is likely responsible for the decreased rate of elemental release from the rock matrix. When the CA1 gene was functional, all organisms, including the WT, CA2, and CA3, overcame the rock’s buffering capacity and acidified the 1% w/v basalt media to pH 5.11, 5.43, and 4.24, respectively. Interestingly, in Figure 3b there were no significant differences in dissolved inorganic carbon seen between organisms, suggesting that the loss in weathering potential was not due to the reduction in carbonic acid catalysis. However, as pH determines carbonate speciation in solution (Wang et al., 2013), with dissolved CO_2_ and carbonic acid predominant at a pH < 6, bicarbonate between pH 6 and 10, and carbonate at pH > 10, this explains the lack of difference observed. As the pH values of the wildtype, CA2, and CA3 media were all < 6, they likely lost dissolved inorganic carbon as gaseous CO_2_ during the experiment. In contrast, the CA1 mutant, which had a pH of 7.11, likely retained inorganic carbon in the dissolved bicarbonate form. Therefore, CA1 inactivation likely resulted in reduced rates of carbonic acid formation, thereby decreasing rates of weathering, and preventing the bacteria from overcoming the buffering capacity of the 1% w/v basalt. These effects of pH buffering caused by the basalt concentration are summarized in Figure 6, which sets out hypotheses to explain the findings. However, it is important to note that CA1 inactivation may have affected weathering rates indirectly through altering connected cellular processes. When all three CA genes found within *B. thailandensis* were assessed using BlastKOALA (Kanehisa et al., 2016), an annotation software, and assigned a KEGG Orthology classification number they were all assigned to the K01673 classification, despite having different sequences. Using KEGG’s Pathways tool it predicted that the K01673 group could play a role in broader metabolic pathways, including the nitrogen metabolism, suggesting that CA1 gene inactivation could have indirectly affected weathering processes through altering other downstream cellular processes.

**Figure 6.**
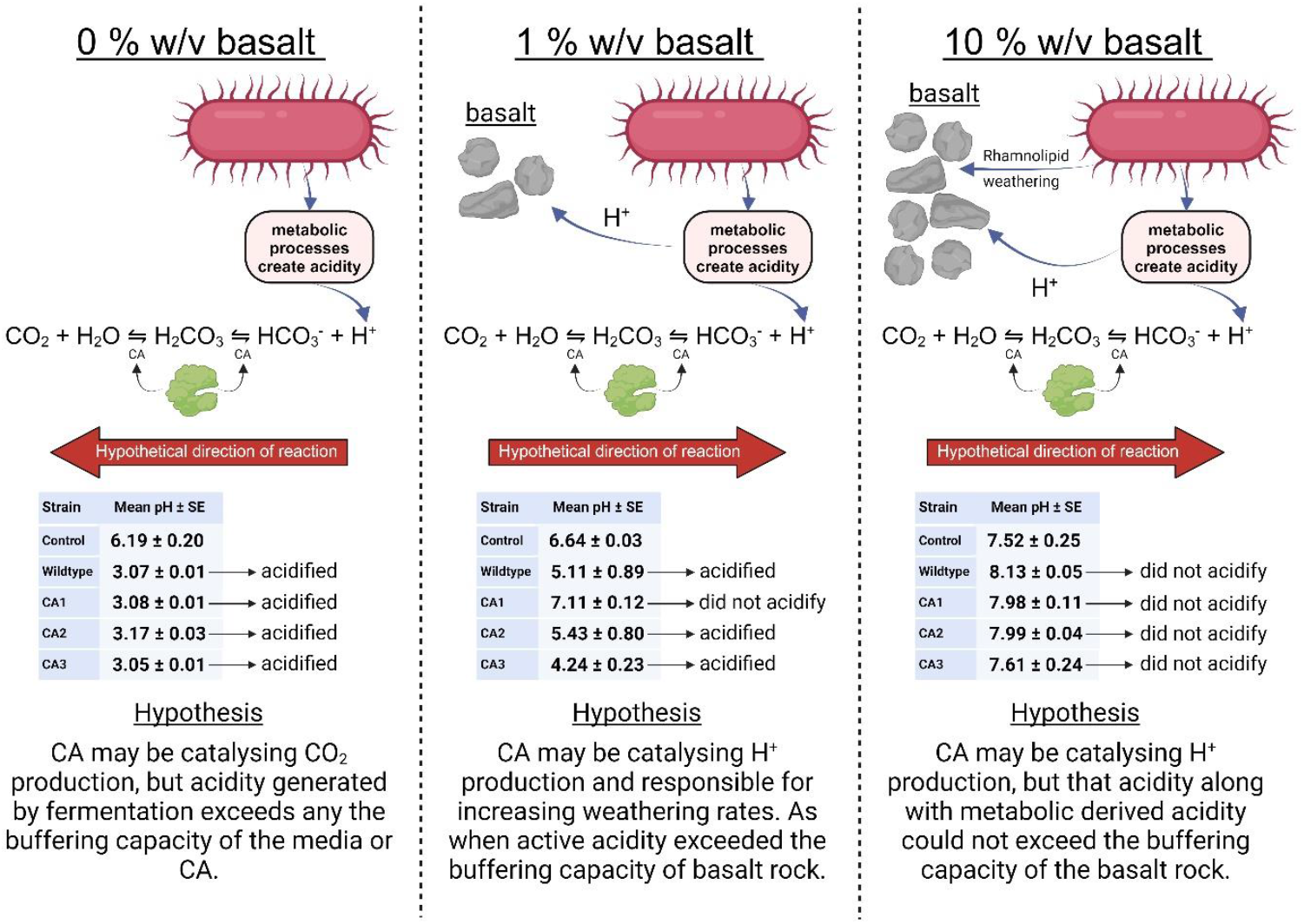
Conceptualisation of CA derived weathering at the three basalt concentrations of 0, 1, and 10%. Created in Biorender.com

Sun and Lian (2019) showed that the two CA genes within *A. nidulans* exhibited distinct physiological functions, with only *canB* being important for acclimation at low ambient CO_2_ concentrations. This is also the case in *B. thailandensis*, which carries three β-CA’s, as the mutant organisms responded differently to basalt amendment. Both the CA2 and CA3 CA genes did not appear to provide redundancy for the CA1 CA gene. These results introduce another layer of complexity to our understanding of CA’s catalytic potential within soils, as not only do CAs between organisms have differing catalytic potentials, but also genes within the same organism carry functional differences. Furthermore through using the cellular localisation tool PSORTb v. 3.0.3 (Yu et al., 2010) all three *B. thailandensis* CA encoding genes were predicted to be intracellularly localised suggesting that intracellular CA’s may play an important role in rock weathering, contradicting previous research that emphasized the importance of extracellular CA. Furthermore, these results may also run contrary to Sauze et al. (2018), who found that bovine CA becomes more efficient at converting CO_2_ to carbonic acid with increasing soil pH at least to a soil pH of 8.5. Our results suggest that CA played a greater role in cation dissolution when *B. thailandensis* was grown in a mildly buffered media (low basalt w/v) compared to highly buffered media (high basalt w/v) with a greater initial pH.

When grown in the 10% basalt media the DECs (Supplemental Figure 1) showed no organism specific significant differences for almost all the measured elements. However, there were non-significant reductions in the elements Ti, V, Al, Co, Ni, Ag, Cs, Fe, and Ba in the CA1 mutant compared to WT. Through performing a three-way ANOVA of the DEC using some of the above elements: there was a significant reduction in the average elemental release compared to the WT and the CA3 mutant. This potential reduction in weathering is not observed when base cation DECs are summed (Figure 5b), suggesting that the inactivation of the CA1 and CA2 genes affects rock weathering differently in the 1% w/v media compared to other conditions, possibly indicating that the CA genes influence cellular processes or behave differently under varying conditions. A further potential explanation for the difference in trends seen between the 1% and 10% w/v basalt media is that *B. thailandensis* produces rhamnolipids. Rhamnolipids, a glycolipidic biosurfactant, which can extract rare earth elements, functions more favourably at a higher pH (Castro et al., 2023).

There was a significant reduction in dissolved phosphorus because of *B. thailandensis* inoculation. *B. thailandensis* is known to carry phosphate solubilising genes (Sivaji et al., 2016) and has been seen in soils to mobilise phosphate (Panhwar et al., 2015). One explanation for the loss in dissolved phosphorus is that as it is required for DNA, RNA, and phospholipid production, and is likely to be assimilated into the bacteria. As no bacterial lysis step was performed the biomass phosphorus was likely lost in the form of pelletised cellular debris during the centrifugation and filtration steps following incubation.

Three E264 B. thailandensis mutants with knoclassificationuts of the genes gluconate kinase (*gnt*K), glucose dehydrogenase (*gdh*FAD), or siderophore synthesis (*pch*D) had their dunite weathering capacity previously assessed relative to the wildtype, as reported in the literature (Epihov, 2018). As seen in Figure 7 below, relative to the wildtype none of the KO’s reduced magnesium, silicon, or iron solubilisation/weathering rates by more than 50%. As expected, the inactivation of the siderophore synthesis gene *pch*D resulted in the greatest reduction in iron levels. In contrast, the inactivation of the *gnt*K gene led to an increase in weathering, potentially due to impairments in the pentose phosphate pathway, which could cause acid buildup (Epihov, 2018). As seen in Figure 7 below, the CA1 inactivation appeared to result in a greater loss of rock weathering potential compared to all other mutants previously tested across the three elements. Again, highlighting the potential importance of CA to bacterial derived rock weathering.

**Figure 7.**
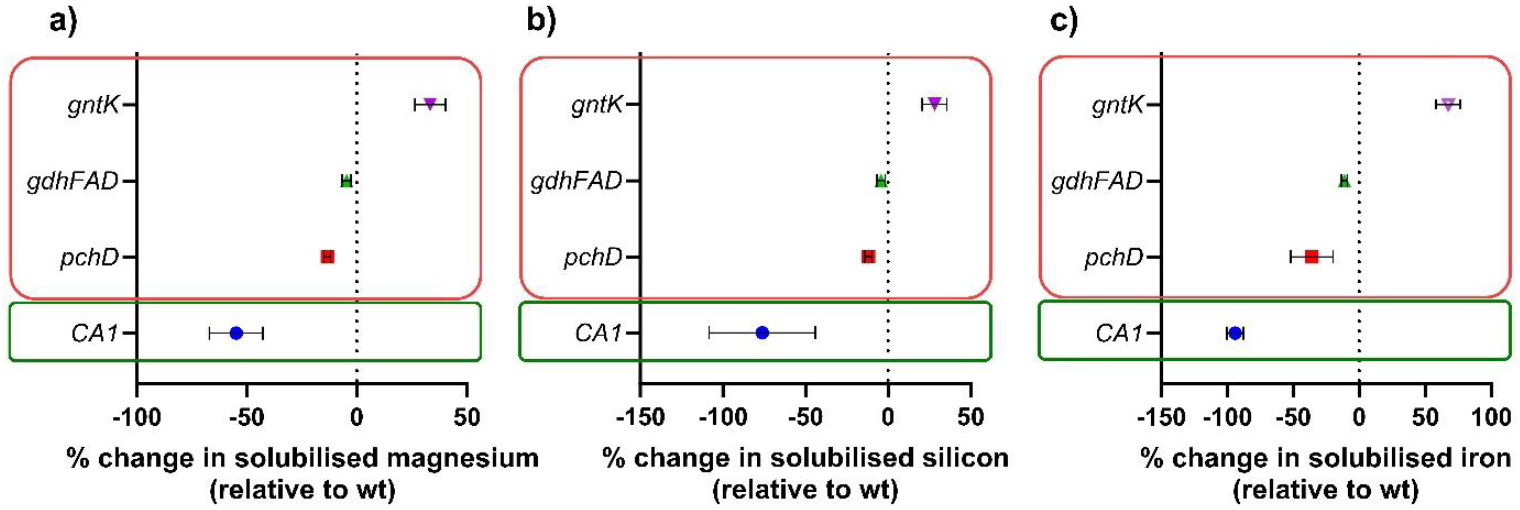
Percent change of solubilised magnesium (a), silicon (b), and iron (c) relative to the wildtype as a result of inactivation of *B. thailandensis* genes gluconate kinase (*gnt*K), glucose dehydrogenase (*gdh*FAD), and siderophore (*pch*D) when incubated with dunite rock, as noted in red and extracted from Epihov (2018), and the CA1 mutant (BTH_I1052) when incubated with basalt rock as seen in green.

Further research is required to establish the extent to which CA can facilitate commercial applications of ERW to farmland and improve the rates of weathering in a cost-effective way. The CDR cost of deploying ERW in the midwestern United States was estimated to be US $45-472 per tonne CO_2_e absorbed, which is comparable to other nature based CDR strategies such as bioenergy with carbon capture and storage, afforestation/reforestation, and biochar (Zhang et al., 2023). Beerling et al. (2020) also calculated CDR costs across farmland in the United States and estimated that by 2050 the cost could range from US $80–180 per tonne of CO_2_ removed. However, improving ERW rates within soils using biotechnology could help improve CDR efficiency by ERW and its technically viability (Epihov et al., 2024). The field of optimising ERW’s CDR rates in soils is nascent, but research has already highlighted its potential. For instance, Epihov et al. (2024) has estimated that the addition of potassium-EDDHA, an iron chelator that may stimulate microbial siderophore production and mineral dissolution, could reduce the CDR costs per hectare to $77 per tonne CO_2_. Whether CA applications can deliver similar cost-benefits remains an open question.

## 5. Conclusion

Our reverse genetic approach indicates that *B. thailandensis* CA influences basalt dissolution rates *in vitro* under mild and highly buffered conditions. However, only one of the three CAs encoded in *B. thailandensis* consistently affected pH and DECs, indicating that bacterial CAs belowground may not all be relevant for rock weathering. The observed reduction in rock weathering upon inactivation of the BTH_I1052 (CA1) CA gene is likely attributable to the KO maintaining a higher pH relative to the WT and other mutants. The increase in solution acidity and rock weathering products following incubation with the other organisms may have been a direct result of carbonic acid formation as concentrations of dissolved inorganic carbon were likely affected by the pH of the solution. Furthermore, when *B. thailandensis* was grown in the highly buffered media, at a pH > 7.5, the BTH_I1052 (CA1) and the BTH_I0345 (CA2) genes do appear to play a role in rock dissolution as the mean DEC was significantly lower than the wildtype and BTH_I1199 (CA3) KO when the solution concentrations of the elements titanium, vanadium, cobalt, nickel, silver, and caesium were averaged. These reductions in weathering were not however noted in cations. Interestingly, CA did not appear to be more functional at the higher pH as larger differences were seen between the wildtype and the KO when grown in the 1% w/v media, indicating pH and buffering capacity as important regulators for the utility of bacterial CAs for ERW-CDR. We conclude that despite its predicted intracellular localization CAs encoded for in bacteria can affect weathering rates and should be considered in the future as a potential target for biologically accelerated ERW in soils.

## 6. CRediT authorship contribution statement

**Derek S Bell:** Conceptualisation, Methodology, Validation, Formal analysis, Investigation, Resources, Data Curation. Writing – original draft, Writing-review & editing, Visualization, Project administration; **Jonathan R. Leake:** Conceptualisation, Methodology, Writing – review & editing, Resources, Supervision, Project administration, Funding acquisition; **David J Beerling**: Validation, Writing – review & editing; **Jurriaan Ton:** Conceptualisation, Writing – review & editing, **Dimitar Z. Epihov** Conceptualisation, Methodology, Validation, Writing – Review & Editing, Supervision, Project administration, Funding acquisition.

## 9. Data statement

Data will be uploaded prior to journal submission.

## 10. Declaration of generative AI and AI-assisted technologies in the writing process

During the preparation of this work, the author(s) used ChatGPT, an AI language model, to assist with grammar and refining content. After using this tool, the author(s) reviewed and edited the content as needed and take(s) full responsibility for the content of the published article.

## 1. Supplementary figures

**Figure 1.**
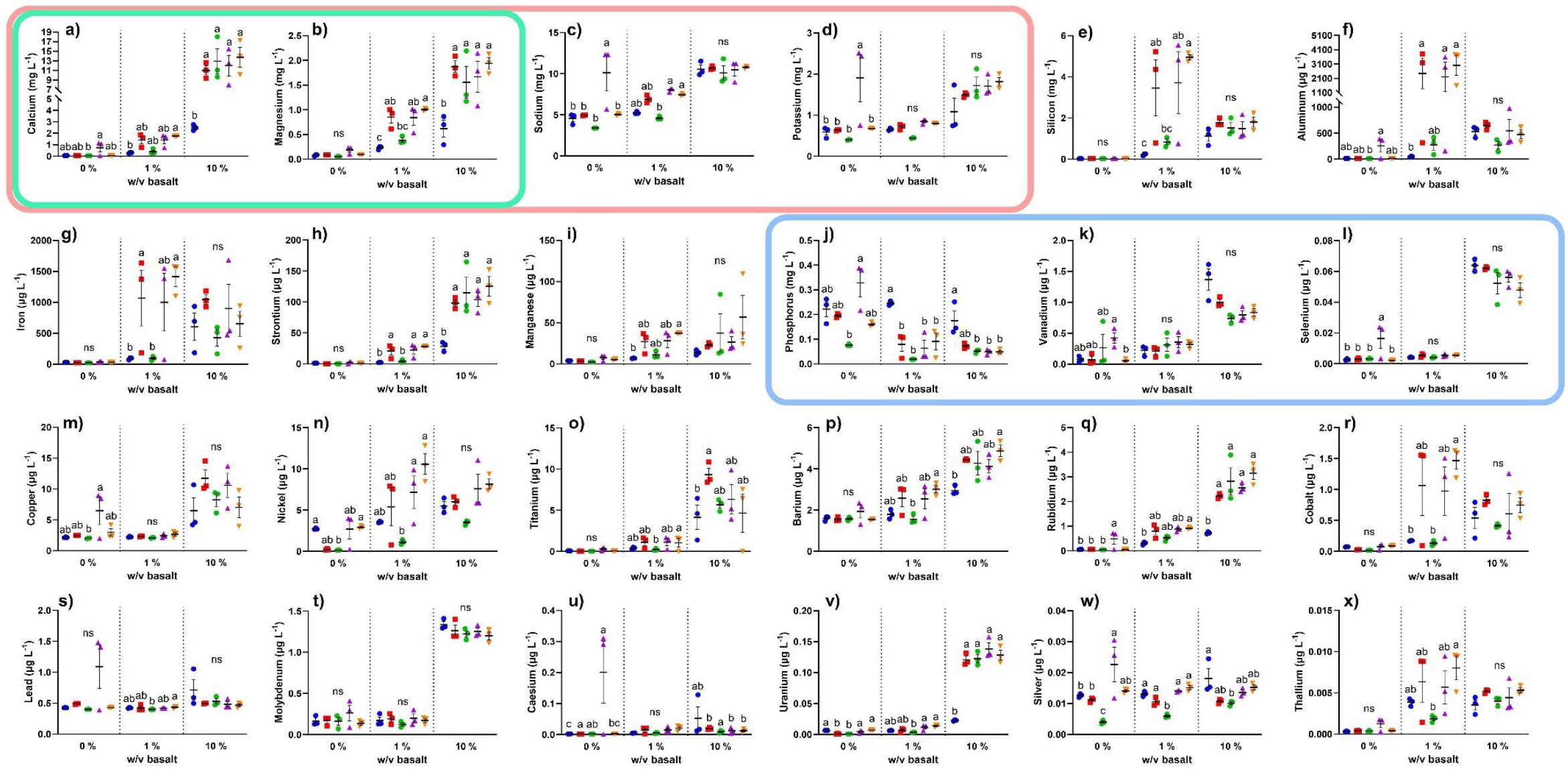
Effects of basalt treatments and *B. thailandensis* Organisms 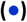 Control, 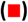 Wildtype, 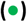 CA1, 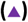 CA2, and 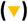 CA3 after 170 hours post inoculation compared to non-inoculated controls on mean (± 1 standard error) dissolved elemental concentrations of (a) calcium, (b) magnesium, (c) sodium, (d) potassium, (e) silicon, (f) aluminium, (g) iron, (h) strontium, (i) manganese, (j) phosphorus, (k) vanadium, (l) selenium, (m) copper, (n) nickel, (o) titanium, (p) barium, (q) rubidium, (r) cobalt, (s) lead, (t) molybdenum, (u) caesium, (v) uranium, (w) silver, and (x) thallium. With dissolved cation ANOVA post hoc Tukey’s test significance differences denoted by letters codes within basalt treatments, where means sharing the same letter are not significantly different (*p* > 0.05). Base cations, calcium and magnesium cations used for calculating potential CDR, and reduced DEC with bacterial inoculation are highlighted by pink, green, and blue frames, respectively.

**Supplemental Table 1.** Two-way ANOVA results of *B. thailandensis* growth media after 170 hours following growth.

| Element | Two-way ANOVA effects | df | F value | P value |
| --- | --- | --- | --- | --- |
| Sodium | w/v basalt | (1, 35) | 142.31 | < 0.0001 |
|  | Organism | (4, 35) | 12.42 | < 0.0001 |
|  | Interaction | (4, 35) | 5.93 | < 0.005 |
| Magnesium | w/v basalt | (1, 35) | 85.96 | < 0.0001 |
|  | Organism | (4, 35) | 4.28 | < 0.05 |
|  | Interaction | (4, 35) | 1.04 | > 0.05 |
| Calcium | w/v basalt | (1, 35) | 273.64 | < 0.001 |
|  | Organism | (4, 35) | 6.37 | < 0.0001 |
|  | Interaction | (4, 35) | 3.26 | < 0.05 |
| Silicon | w/v basalt | (1, 35) | 13.60 | < 0.001 |
|  | Organism | (4, 35) | 0.77 | > 0.05 |
|  | Interaction | (4, 35) | 0.10 | > 0.05 |
| Aluminum | w/v basalt | (1, 35) | 9.24 | < 0.01 |
|  | Organism | (4, 35) | 1.68 | > 0.05 |
|  | Interaction | (4, 35) | 0.62 | > 0.05 |
| Manganese | w/v basalt | (1, 35) | 22.35 | < 0.0001 |
|  | Organism | (4, 35) | 2.80 | < 0.05 |
|  | Interaction | (4, 35) | 0.42 | > 0.05 |
| Iron | w/v basalt | (1, 35) | 24.39 | < 0.0001 |
|  | Organism | (4, 35) | 1.39 | > 0.05 |
|  | Interaction | (4, 35) | 0.20 | > 0.05 |
| Cobalt | w/v basalt | (1, 35) | 16.92 | < 0.001 |
|  | Organism | (4, 35) | 2.08 | > 0.05 |
|  | Interaction | (4, 35) | 0.58 | > 0.05 |
| Strontium | w/v basalt | (1, 35) | 190.30 | < 0.0001 |
|  | Organism | (4, 35) | 4.77 | < 0.01 |
|  | Interaction | (4, 35) | 1.86 | > 0.05 |
| Molybdenum | w/v basalt | (1, 35) | 975.42 | < 0.0001 |
|  | Organism | (4, 35) | 1.00 | > 0.05 |
|  | Interaction | (4, 35) | 0.52 | > 0.05 |
| Silver | w/v basalt | (1, 35) | 7.39 | < 0.05 |
|  | Organism | (4, 35) | 36.09 | < 0.0001 |
|  | Interaction | (4, 35) | 8.12 | < 0.0001 |
| Caesium | w/v basalt | (1, 35) | 0.89 | > 0.05 |
|  | Organism | (4, 35) | 9.19 | < 0.001 |
|  | Interaction | (4, 35) | 3.08 | < 0.05 |
| Phosphorus | w/v basalt | (1, 35) | 8.64 | < 0.001 |
|  | Organism | (4, 35) | 7.03 | < 0.01 |
|  | Interaction | (4, 35) | 1.21 | > 0.05 |
| Potassium | w/v basalt | (1, 35) | 79.50 | < 0.0001 |
|  | Organism | (4, 35) | 8.22 | < 0.001 |
|  | Interaction | (4, 35) | 4.27 | < 0.01 |
| Titanium | w/v basalt | (1, 35) | 242.69 | < 0.0001 |
|  | Organism | (4, 35) | 6.37 | < 0.05 |
|  | Interaction | (4, 35) | 1.41 | > 0.05 |
| Vanadium | w/v basalt | (1, 35) | 148.583 | < 0.0001 |
|  | Organism | (4, 35) | 1.52 | > 0.05 |
|  | Interaction | (4, 35) | 4.97 | < 0.01 |
| Lead | w/v basalt | (1, 35) | 5.21 | < 0.05 |
|  | Organism | (4, 35) | 1.27 | > 0.05 |
|  | Interaction | (4, 35) | 0.11 | > 0.05 |
| Nickel | w/v basalt | (1, 35) | 17.74 | < 0.001 |
|  | Organism | (4, 35) | 6.99 | < 0.001 |
|  | Interaction | (4, 35) | 0.65 | > 0.05 |
| Copper | w/v basalt | (1, 35) | 108.07 | < 0.0001 |
|  | Organism | (4, 35) | 2.66 | < 0.05 |
|  | Interaction | (4, 35) | 1.22 | > 0.05 |
| Uranium | w/v basalt | (1, 35) | 819.37 | < 0.0001 |
|  | Organism | (4, 35) | 13.74 | < 0.0001 |
|  | Interaction | (4, 35) | 23.96 | < 0.0001 |
| Selenium | w/v basalt | (1, 35) | 518.85 | < 0.0001 |
|  | Organism | (4, 35) | 3.38 | < 0.05 |
|  | Interaction | (4, 35) | 2.35 | > 0.05 |
| Rubidium | w/v basalt | (1, 35) | 154.87 | < 0.0001 |
|  | Organism | (4, 35) | 5.87 | < 0.01 |
|  | Interaction | (4, 35) | 3.14 | < 0.05 |
| Barium | w/v basalt | (1, 35) | 135.667 | < 0.0001 |
|  | Organism | (4, 35) | 3.94 | < 0.01 |
|  | Interaction | (4, 35) | 1.43 | > 0.05 |
| Thallium | w/v basalt | (1, 35) | 12.68 | < 0.01 |
|  | Organism | (4, 35) | 0.80 | > 0.05 |
|  | Interaction | (4, 35) | 0.19 | > 0.05 |

